# Analyses of bent spindles reveal the mechanics of anaphase B in fission yeast

**DOI:** 10.1101/2025.11.20.689516

**Authors:** Paula Real-Calderón, Thomas G. Fai, Rafael R. Daga, Joël Lemière, Fred Chang

**Affiliations:** Department of Cell and Tissue Biology, University of California San Francisco, San Francisco, CA 94117, United States; Department of Nuclear Architecture and Dynamics, Andalusian Center for Development Biology (CABD), Seville, Spain, 41013; Department of Mathematics and Volen Center for Complex Systems, Brandeis University, Waltham, MA 02453, United States

## Abstract

The mitotic spindle in the fission yeast *Schizosaccharomyces pombe* is a single bundle of microtubules which elongates to segregate the chromosomes during anaphase B. The mechanical properties of the spindle and the forces driving its elongation remain poorly defined. Here, we analyzed how spindles react to mechanical and genetic perturbations to uncover their mechanical properties. Treatment of cells with osmotic oscillations and blue light led to a consistent phenotype of spindle buckling and breakage in mid-anaphase. The stalling of pole separation and reduced rates of spindle elongation indicated that spindles elongate and buckle under increased mechanical load. The structural integrity of the bent spindles was dependent on Ase1 (PRC1), while the spindle elongation rate was dependent on motor proteins Klp9 (kinesin-6) and Cut7 (kinesin-5). Modeling of bent spindle shapes revealed that most spindles behave mechanically as a beam with a two-fold increase in rigidity in the midzone. Upon reaching a threshold size, bent spindles broke at a specific fragile site near the edge of the spindle midzone. Our findings in this simple fission yeast spindle are relevant to the mechanics of more complex metazoan spindles.

**Significance statement:**

- The anaphase B spindle in *S. pombe* consists of a microtubule bundle that elongates to move the chromosomes apart. The various forces and mechanical properties of the spindle remain poorly quantified.
- The authors establish a method to induce spindle buckling in mid-anaphase. Time-lapse imaging shows that these spindles elongate at reduced rates, buckle as a non-homogeneous beam under mechanical load, and break at a fragile site adjacent to the midzone.
- These results provide quantitative and molecular insights into spindle force regulation and structural integrity that are relevant to mitosis in other cell types.

## Introduction

Mitosis involves the large-scale reorganization and movement of cellular components, driven by internal cellular forces. While molecular players involved in mitotic spindle assembly and chromosome segregation have been characterized (Edelmaier et al., 2020), the physical principles governing force generation and the material properties of the spindle remain poorly understood. Quantifying spindle forces poses a significant challenge due to the spindle’s small size, its complex and dynamic behavior, and its location within the cell. One of the first measurements of spindle forces was conducted by Nicklas, who employed glass microneedles to probe the exceptionally large spindle in grasshopper spermatocytes (Nicklas, 1983). Estimates of spindle forces by various approaches however range over two orders of magnitude (Nicklas, 1983; Jannink et al., 1996; Marshall et al., 2001; Ferraro-Gideon et al., 2013). To integrate our understanding of spindle mechanics and molecular-based mechanisms across various cell types, the development of additional non-invasive approaches for measuring and manipulating spindle forces is needed.

The fission yeast *Schizosaccharomyces pombe* provides a simple model system to study the eukaryotic spindle. Its spindle is composed of a single antiparallel bundle of microtubules (MTs), which are nucleated from the spindle pole bodies (SPBs) embedded in the nuclear envelope (NE) (Ding et al., 1993; Ward et al., 2014). Mitosis occurs in discrete stages in a highly stereotypical manner: upon chromosome capture and alignment at metaphase, the kinetochores move towards the SPBs during anaphase A, followed by movement of the SPBs and chromosomes towards the cell poles in anaphase B (Nabeshima et al., 1998). The elongation of the spindle in anaphase B is primarily driven by MT sliding and MT plus-end polymerization at the spindle midzone, in the absence of MT minus-end flux at the SPBs (Mallavarapu et al., 1999; Khodjakov et al., 2004; Lera-Ramirez et al., 2022). Unlike many eukaryotic systems, cytoplasmic astral MTs are dispensable for this process (Tolic-Nørrelykke et al., 2004; Zimmerman & Chang, 2005).

A set of conserved molecules required for spindle formation and elongation in fission yeast has been functionally characterized through genetic interactions, live cell imaging, and *in vitro* studies (Glotzer, 2009; Salas-Pino & Daga, 2019; Anjur-Dietrich et al., 2021; Vukušić & Tolić, 2021). Klp9 (kinesin-6) is the best candidate for the primary spindle anaphase B motor as well as a MT polymerase (Yukawa et al., 2019; Krüger et al., 2021). Other spindle motor proteins include the plus-end directed motor Cut7 (kinesin-5) and the minus-end directed motors Klp2 and Pkl1 (both kinesin-14s) (Yukawa et al., 2019; Loncar et al., 2020; Krüger et al., 2021). Besides these motors, other spindle proteins such as XMAP215/Alp14/Dis1 (Garcia et al., 2001; Yukawa, Kawakami, et al., 2019), Cls1/Peg1/CLASP (Bratman & Chang, 2007) and Ase1 (PRC1/MAP65) (Loiodice et al., 2005) all contribute in different ways to spindle integrity and elongation. Notably, Ase1 crosslinks antiparallel MTs at the midzone, where it recruits proteins such as Klp9 and Cls1/Peg1 (CLASP) (Bratman & Chang, 2007; Fu et al., 2009). Collectively, these studies have defined many of the key parameters and molecular factors necessary for quantitative models of spindle mechanics (Ward et al., 2014; Lamson et al., 2019; Edelmaier et al., 2020).

Here, we introduce an approach to probe the mechanical properties of the spindle *in vivo* by quantitative analyses of spindle buckling. In previous papers, our group has used osmotic oscillations in fission yeast cells to investigate mechanical aspects of cell growth and MT dynamics (Knapp et al., 2019; Molines et al., 2022). For example, during hyperosmotic shocks, cells exhibit a rapid, reversible decrease in cellular and nuclear volume (up to 50%), increased cytoplasmic concentration, and reduced rates of MT growth and shrinkage (Knapp et al., 2019; Lemière et al., 2022; Molines et al., 2022; Lemière & Chang, 2023).

In this study, we build upon the observation that fission yeast cells exposed to osmotic oscillations exhibit a dramatic failure in anaphase -- not during the osmotic oscillations, but 1-2 hr later in the subsequent mitosis. These cells exhibited a surprisingly consistent mitotic phenotype, in which spindles elongate and bend during anaphase, and then abruptly break into two fragments at a defined fragile site. This large deformation resembles the behavior of a beam that buckles and breaks spontaneously under mechanical load. By analyzing the elongation rates and geometric shape of these bent spindles, we derive quantitative insights into spindle elongation forces and its mechanical properties under load.

## Results

### Transient treatment of osmotic oscillations produces anaphase failure in the subsequent mitosis

This study was inspired by an unexpected observation: treating fission yeast cells with osmotic oscillations produced mitotic failures over an hour after the oscillations had ended. In a typical experiment, yeast cells were grown in a microfluidic device and imaged using a spinning disc confocal microscope with lasers at 488 nm and 561 nm. Cells were first grown in rich media (YES), then treated with hyperosmotic shocks in an oscillatory pattern (alternating between YES and YES+1 M sorbitol every 5 min, repeated five times), and then returned to YES. After the oscillations, cells promptly resumed cell growth, often dividing and proceeding through interphase of the next cell cycle (Knapp et al., 2019). However, when these cells entered mitosis again, over 70% exhibited mitotic failures during anaphase (Figure 1A and 1B), producing the classic “cut” phenotype in which cells attempted to septate without a proper nuclear division (Hirano et al., 1986; Funabiki et al., 1996).

**Figure 1.**
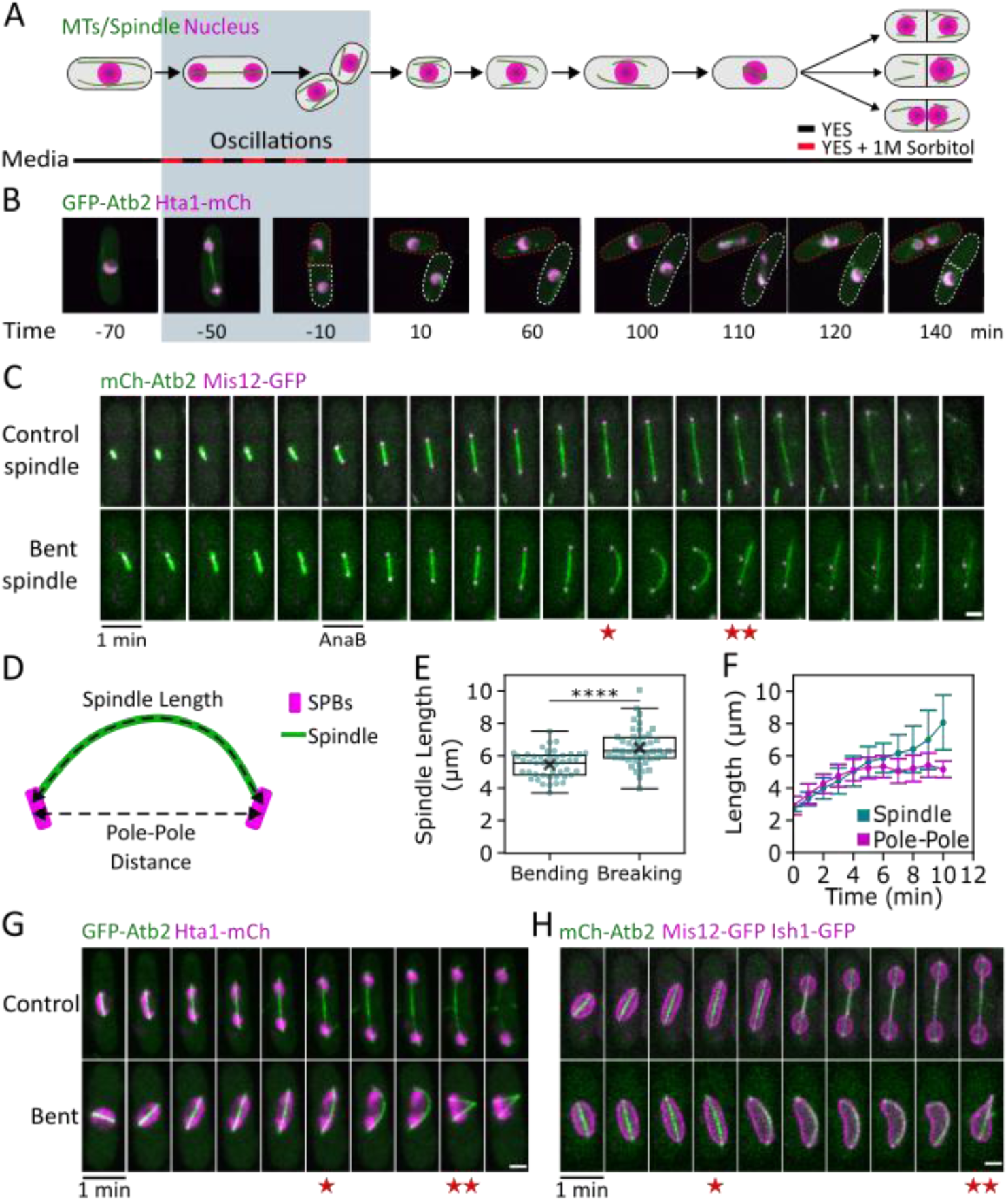
Osmotic oscillations produce anaphase failure in the subsequent mitosis. **(A)** Schematic of the protocol to induce anaphase failures in fission yeast cells. MTs (green) and nuclei (magenta) are depicted. **(B)** Maximum intensity projection confocal images of a WT cell expressing a marker for MTs (tubulin GFP-Atb2, in green) and for chromosomes (histone Hta 1-mCh, magenta) treated with the protocol described in (A). The cell divides normally during the osmotic oscillations (t = −50 to 0 min), but subsequently, its daughter cells growing in normal media exhibit anaphase defects 1-2 hr later (t = 60-120 min). **(C)** Time-lapse images of MTs (mCh-Atb2, green) and kinetochores (Mis12-GFP, magenta) in a cell undergoing mitosis in control conditions with no osmotic treatment (control) and in a cell after the osmotic treatment that exhibits a spindle that bends and breaks (bent). Anaphase B onset is indicated with Ana B. Stars indicate points of spindle bending (★) and breaking (★★). **(D)** Schematic of how the spindle length and the pole-to-pole distances were measured. **(E)** Spindle lengths at initiation of bending and breaking (n = 45). **(F)** Mean spindle length and pole-to-pole distances over time. t = 0 corresponds to anaphase B onset; the last time point corresponds with the spindle breakage (n = 29). **(G, H)** Time-lapse images of control cells (top) and cells with bent spindles (bottom), showing behavior of (G) histones (Hta 1-mCh, magenta) and MTs (GFP-Atb2, green); and (H) NE (Ish1-GFP, magenta), kinetochores (Mis12-GFP, magenta), and MTs (mCh-Atb2, green). Scale bar = 2 μm

**Supplemental Figure S1.**
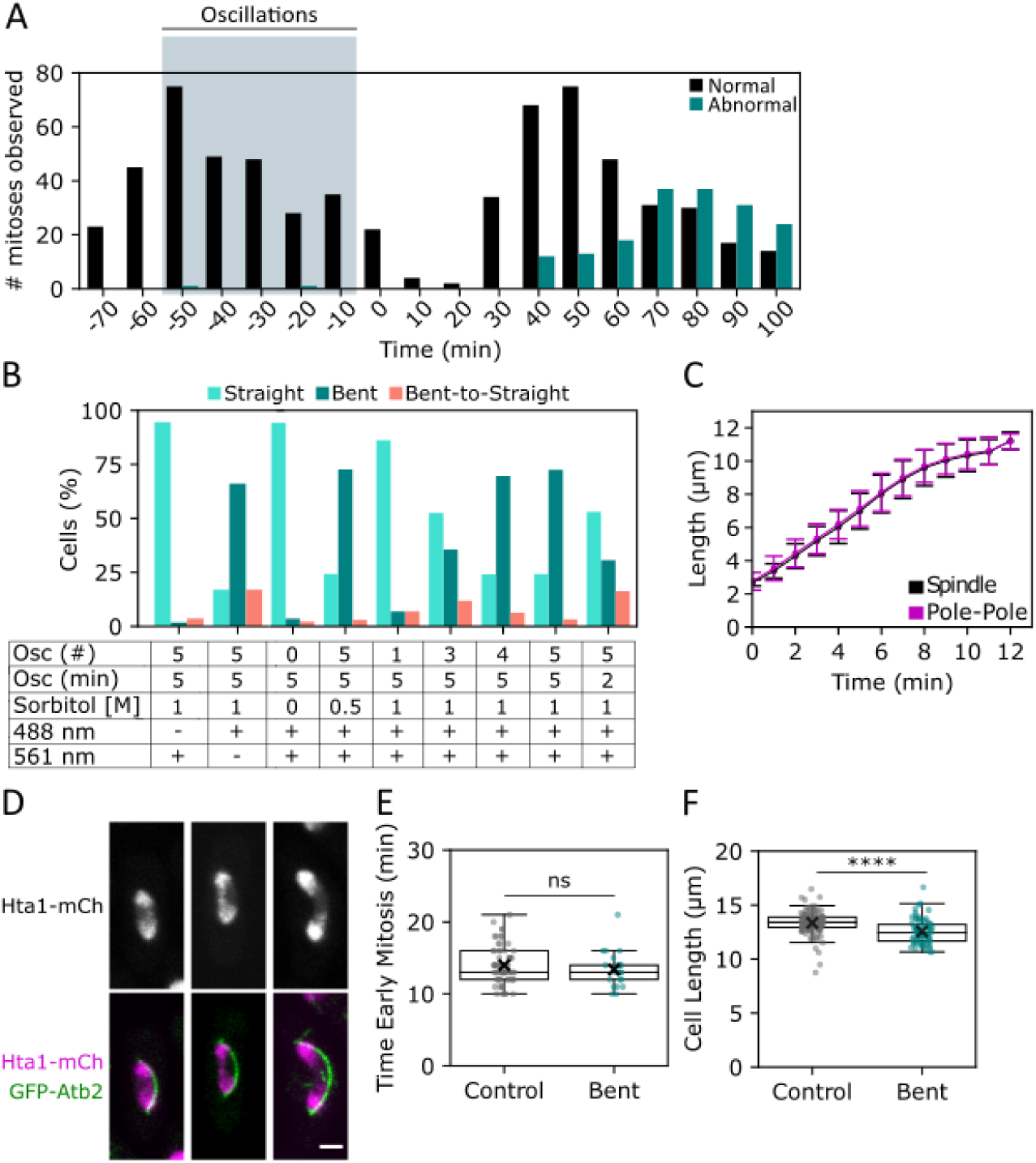
Multiple osmotic oscillations with blue light during interphase produce anaphase failure without triggering DNA damage or spindle cell cycle checkpoints. **(A)** An asynchronous WT cell population, with tubulin (mCh-Atb2) and kinetochores (Mis12-GFP) tagged, was treated with our protocol of osmotic oscillations (see Methods and Figure 1A) and then monitored in time-lapse movies for mitoses. For the graph, mitotic events were classified as normal or abnormal at a single time point at the end of mitosis (n = 822 total). **(B)** Experimental parameters required for the bent spindle phenotype. Cells were treated under different conditions and assayed for mitotic defects in subsequent mitoses. Cell phenotypes were categorized as follows: i) straight spindle that did not break (Straight); ii) bent spindle that eventually broke (Bent); iii) bent spindle that recovered to straight without breaking (Bent-to-Straight). The first two columns show the effects of 488 nm and 561 nm laser light, used every 5 min, during confocalspinning disc imaging (n = 55/59; see Methods). The other columns show the effects of osmotic oscillation number (Osc #), oscillation period (min), and concentration of sorbitol ([M]) during the osmotic oscillations (n = 139/33/29/59/79/182/49). **(C)** Mean spindle length and pole-to-pole distances (µm) in WT control cells with MTs (mCh-Atb2) and kinetochores (Mis12-GFP) tagged. t = 0 corresponds with the onset of anaphase B; the last point corresponds with the disassembly of the anaphase spindle (n = 100). **(D)** Additional examples of anaphase bridges in bent spindles just before spindle breakage (Figure 1G). Scale bar = 2 μm. **(E)** To determine if the treatment triggered the spindle checkpoint in early mitosis, the time in early mitosis (until anaphase B onset) was measured in time-lapse images (n = 64/23). **(F)** To test if the treatment triggers a DNA damage checkpoint, we measured cell size at mitotic entry. Activation of the DNA damage checkpoint causes a cell cycle delay at the G2/M phase, leading to an increase in cell size (Furnari et al., 1997). Cells with bent spindles (green) (n = 74) did not show a larger mitotic cell length than control cells (grey) (n = 117). Instead, they showed a slight reduction in cell length, possibly as a consequence of osmotic treatments.

We were struck by the fact that this treatment did not immediately cause mitotic failure; instead, the effects were delayed by ∼1-2 hr, first appearing in the population around 40 min after the oscillations ended, with the peak percentage of abnormal mitoses occurring 70-90 min later. (Figure 1A and 1B; Supplemental Figure S1A). This delay suggested that cells may be particularly susceptible to the osmotic treatment and blue light during a specific phase(s) of the cell cycle. To test this, we followed individual cells in time-lapse imaging. Cells already in mitosis during the oscillations successfully completed division normally (100%), whereas the majority of cells in interphase during the oscillations developed bent spindles in the subsequent mitosis (80%). Such delayed effects are reminiscent of reports where perturbations experienced during interphase or the previous mitosis result in mitotic catastrophe (Sazonova et al., 2021; Saxena & Zou, 2022).

In testing which experimental perturbations are relevant to produce this phenotype, we found that multiple (3 to 4) osmotic oscillations and exposure to 488 nm blue light (but not 561 nm light) during the oscillations are necessary (Supplemental Figure S1B). Thus, these findings defined a novel non-genetic approach to induce anaphase failure robustly.

### Mitotic failures are caused by spindle bending and breakage

To characterize this mitotic defect, we imaged cells expressing the tubulin marker mCh-Atb2 (Tatebe et al., 2001) and the kinetochore marker Mis12-GFP (Goshima et al., 1999) (Figure 1C). In control cells, the anaphase spindle appeared as a straight MT bundle that elongated steadily from 2.5 to ∼11 µm in length, pushing the spindle poles and kinetochores to the two opposite ends of the cell (Figure 1C and 1D, Supplemental Figure S1C) (Nabeshima et al., 1998). In cells treated with osmotic oscillations and blue light, the early stages of mitosis (prophase and metaphase) proceeded apparently normally. In anaphase A, kinetochores marked by Mis12-GFP moved normally from the metaphase plate to the spindle pole. However, during anaphase B, even though spindles began to elongate at the normal length (∼2.5 µm) as a straight bundle, they began to buckle in mid-anaphase at 5.4 ± 0.81 µm. They continued to grow into a “C” shape until they broke at 6.5 ± 1.15 µm in length (Figure 1E). The bent spindles adopted only “C” shapes, indicative of first-order buckling, and not shapes such as an “S”, indicative of higher-order buckling. Pole-to-pole distances initially increased with spindle length but then stalled in mid-anaphase, remaining constant from the onset of bending until spindle breakage (Figure 1F, see also Figure 3A). Thus, spindle buckling occurred because the spindle continues to elongate while its poles are somehow constrained from separating further. These observations suggest a mechanical model in which resistive forces prevent the poles from separating in mid-anaphase, but the spindle continues to exert forces to elongate the MT bundle, exerting sufficient force to buckle and ultimately break itself.

Previous studies have shown that similar anaphase failures may arise from defects in sister chromatid segregation and/or defects in NE expansion during anaphase (Nakazawa et al., 2016; Begley et al., 2025) (see Discussion). To test these possibilities for this case, we imaged the dynamics of the chromosomes and NE in bent spindles using the histone marker Hta1-mCh (Yamamoto et al., 2019) and the NE marker Ish1-GFP (Taricani et al., 2002). At time points just before spindle collapse, although most of the DNA and the kinetochore had segregated towards the poles, the chromosomes had not clearly separated spatially and appeared to be connected by anaphase bridges (Figure 1G, Supplemental Figure S1D). Upon spindle breakage, the chromosomes collapsed back together. These chromosomal behaviors were consistent with a sister chromatid segregation defect.

The NE formed abnormal shapes, wrapping around the spindle and chromosomes and adopting an ovoid bean shape with a concavity on the side opposite to the spindle without ever adopting the typical dumbbell shape characteristic of a normal nuclear division (Figure 1H). No tearing of the NE was seen. Furthermore, we detected no cell cycle delays in G2 or pre-metaphase, which might indicate activation of DNA damage G2 checkpoints or spindle metaphase checkpoint (Supplemental Figure S1E and S1F) (Enoch & Nurse, 1990; Patterson et al., 1997; Rhind & Russell, 1998). Thus, although these initial findings did not yet definitively demonstrate the underlying cause of this phenotype, we could use this highly penetrant and consistent bent spindle phenotype to probe the mechanical properties of the spindle.

### The molecular requirements for bent spindle formation

We next addressed which spindle components contribute to the elongation forces and structural integrity needed to form and maintain the bent spindle. We hypothesized that in mutants with diminished spindle elongation forces, their spindles might stall or fail to bend, while those with compromised structural integrity might collapse under increased mechanical load.

A core group of anaphase B regulators includes MT motor proteins Klp9 (kinesin-6), Cut7 (kinesin-5), and Klp2 (kinesin-14), and the MT crosslinker Ase1 (PRC1) (Yukawa et al., 2018; Yukawa et al., 2019). While *klp9Δ* and *klp2Δ* cells are viable (Troxell et al., 2001; Fu et al., 2009), Cut7 is essential for cell viability; thus, to assess its contributions, we used a *cut7Δpkl1Δ* double mutant, in which the *pkl1* deletion suppresses the essential role of Cut7 in spindle assembly during early mitosis (Pidoux et al., 1996)*. klp9Δ*, *klp2Δ*, and *cut7Δpkl1Δ* mutants all formed bent spindles after osmotic treatment at comparable frequencies and spindle lengths to the WT strain (Figure 2A, 2C, and 2D). As with the WT spindles, 100% of the bent spindles in these mutants broke (Figure 2A and 2C). Thus, none of these motor proteins were individually required for bent spindle formation.

**Figure 2.**
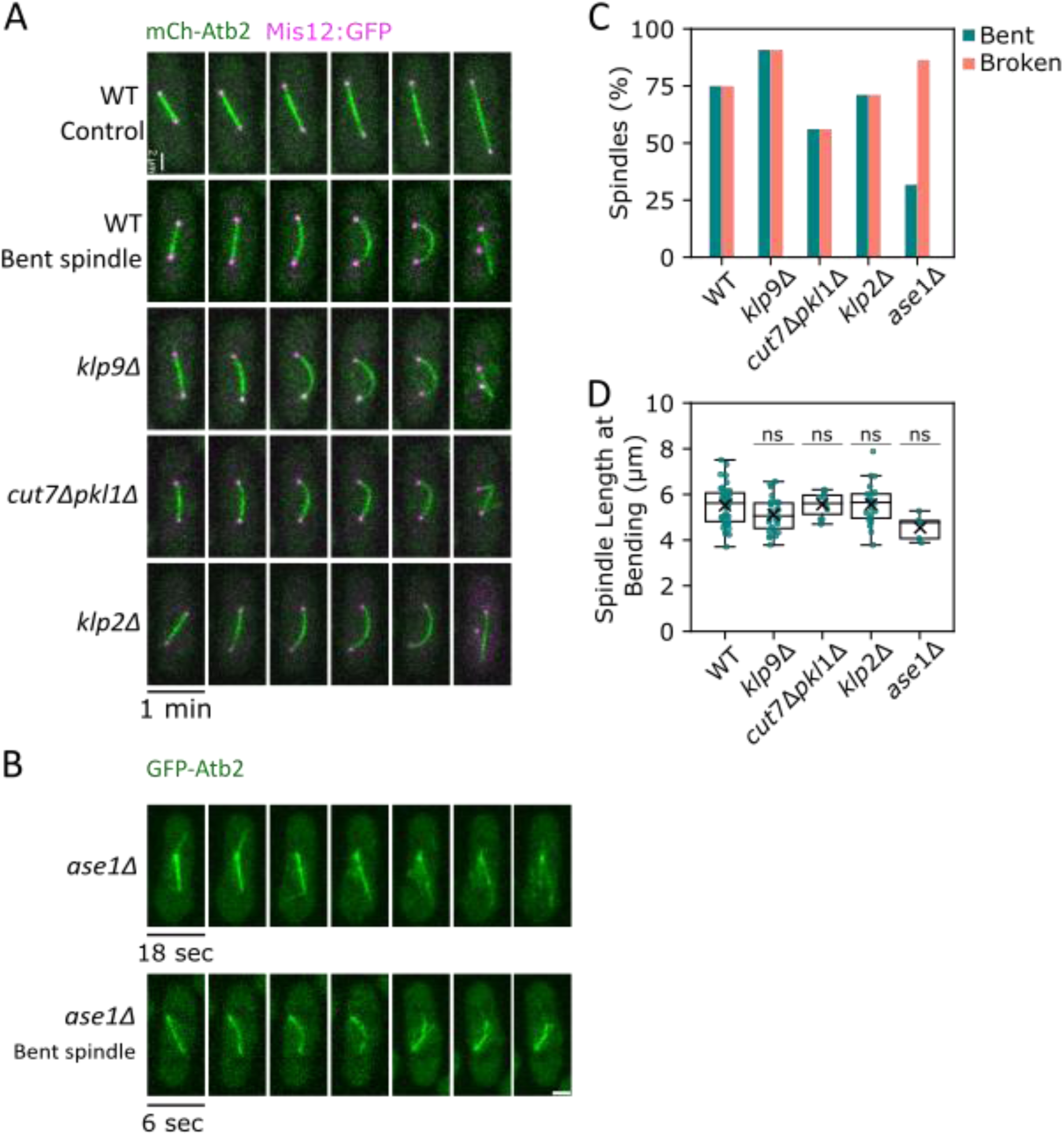
Effects of spindle proteins on spindle bending. **(A)** Time-lapse confocal images of mitotic cells with tagged MTs (mCh-Atb2, green) and kinetochores (Mis12-GFP, magenta). The top panel shows a control WT spindle; the remaining panels depict bent spindles in WT and kinesin-like motor protein mutants after osmotic oscillation treatment. **(B)** Time-lapse images of spindles in *ase1Δ* cells expressing tubulin (GFP-Atb2, green), showing spindle breakage without bending (top) or with transient bending (bottom). Scale bar = 2 µm. **(C)** Percentage of cells treated with the osmotic protocol that showed bent and/or broken spindles. In images of *ase1Δ* spindles taken every 6 s, most spindles broke prematurely without detectable bending (n = 176/66/82/90/22). **(D)** Spindle length at the onset of bending for WT cells and mutant strains. Each dot represents a spindle (n = 30/19/9/21/22).

In contrast, *ase1Δ* mutants, which form normal-appearing spindles without osmotic treatment (Loiodice et al., 2005; Bratman & Chang, 2007), rarely produced stable bent spindles after treatment (Figure 2B and 2C). High-frequency imaging (every 6 sec) revealed that 85% of *ase1Δ* spindles appeared to fall apart at an average spindle size of 5 µm. 63% of these spindles disintegrated into a disorganized array of MTs without any apparent bending, while 37% showed transient bending (most < 30 sec) before breaking (Figure 2B, bottom panel). Therefore, these observations showed that Ase1, but not the individual motor proteins, is needed for the structural integrity of the bent spindle under mechanical load.

### Bent spindles exhibit reduced spindle elongation rates throughout anaphase

To assess the forces responsible for spindle elongation, we compared the rates of anaphase B elongation in WT control and bent spindles (Figure 3A). In fission yeast, for reasons still not completely understood, anaphase B occurs at a near-constant rate (Krüger et al., 2019). Rates of elongation depend on 1) elongation forces from MT sliding and polymerization (Mallavarapu et al., 1999; Krüger et al., 2021), and 2) forces resistive to elongation, such as viscous drag from the cytoplasm and resistive forces from the chromosomes and possibly NE (Ward et al., 2014; Lawrimore & Bloom, 2022; Begley et al., 2025).

**Figure 3.**
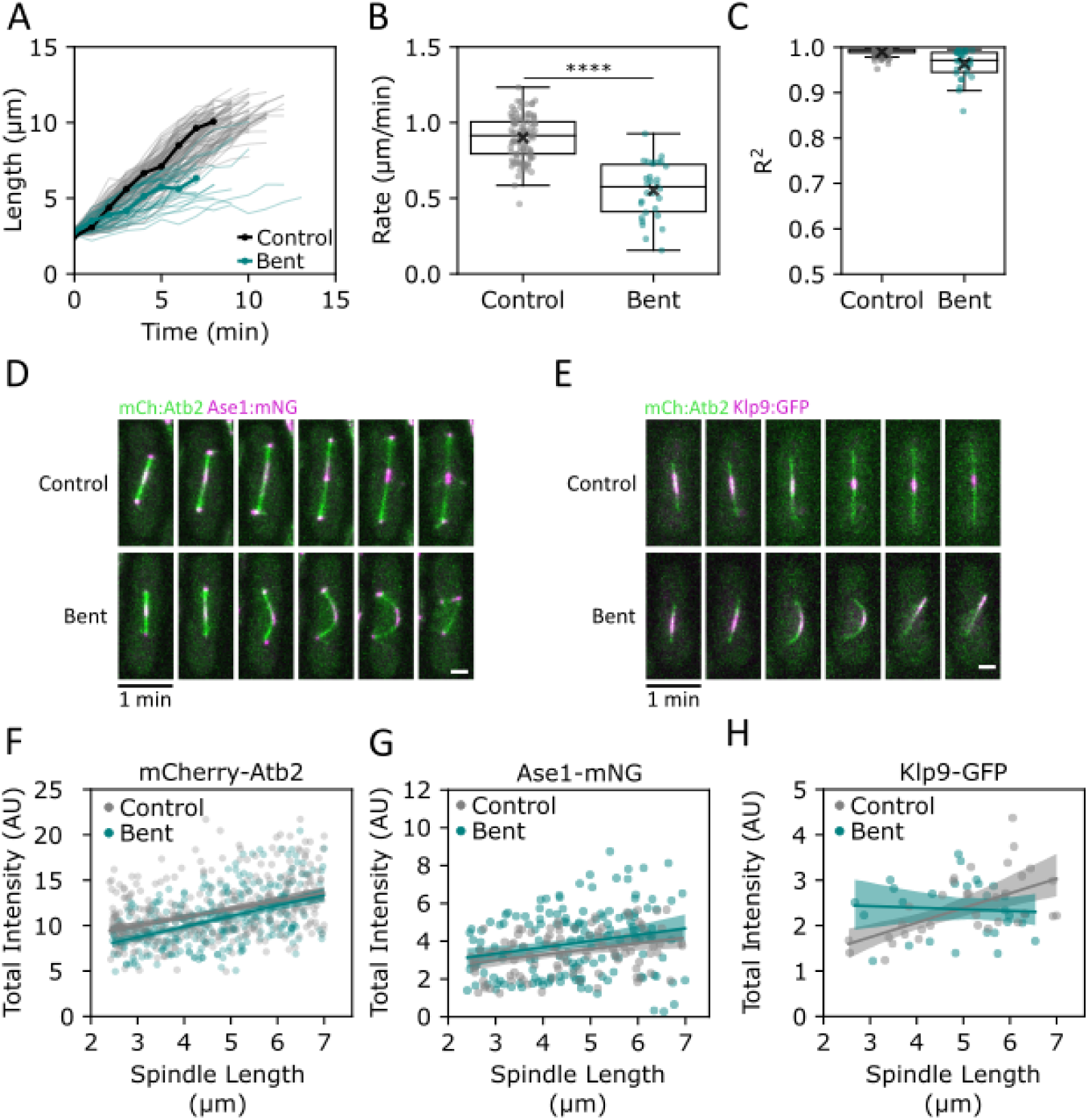
Spindle elongation rates are proportionally decreased throughout anaphase in bent spindles. **(A)** Anaphase B spindle elongation plots for WT cells under control conditions (YES, black) and for bent spindles (green). Each line represents one spindle; representative spindles are highlighted in bold (n = 100/30). **(B, C)** Plot and coefficient determination (R^2^) of the mean elongation rate in control and bent spindles from (A) during anaphase B, assuming a constant elongation rate. **(D)** Confocal maximum intensity projection images _of_ control and bent spindles with mCh-Atb2 (green) and Ase1-mNG (magenta) tagged. **(E)** Same as in (D), but with Klp9-GFP (magenta). Scale bar = 2 µm. **(F)** Summed intensities of mCh-Atb2 in the spindle, plotted as a function of spindle length from 2.5 to 7 µm in control and bent spindles (n = 69/52). **(G, H)** Summed intensities of Ase1-mNG (n = 27/31) and Klp9-GFP (n = 47/22) in the spindle midzone region plotted as a function of spindle length. Each condition was fitted using a linear regression model (line) with CI = 95% (shaded areas).

**Supplemental Figure S3.**
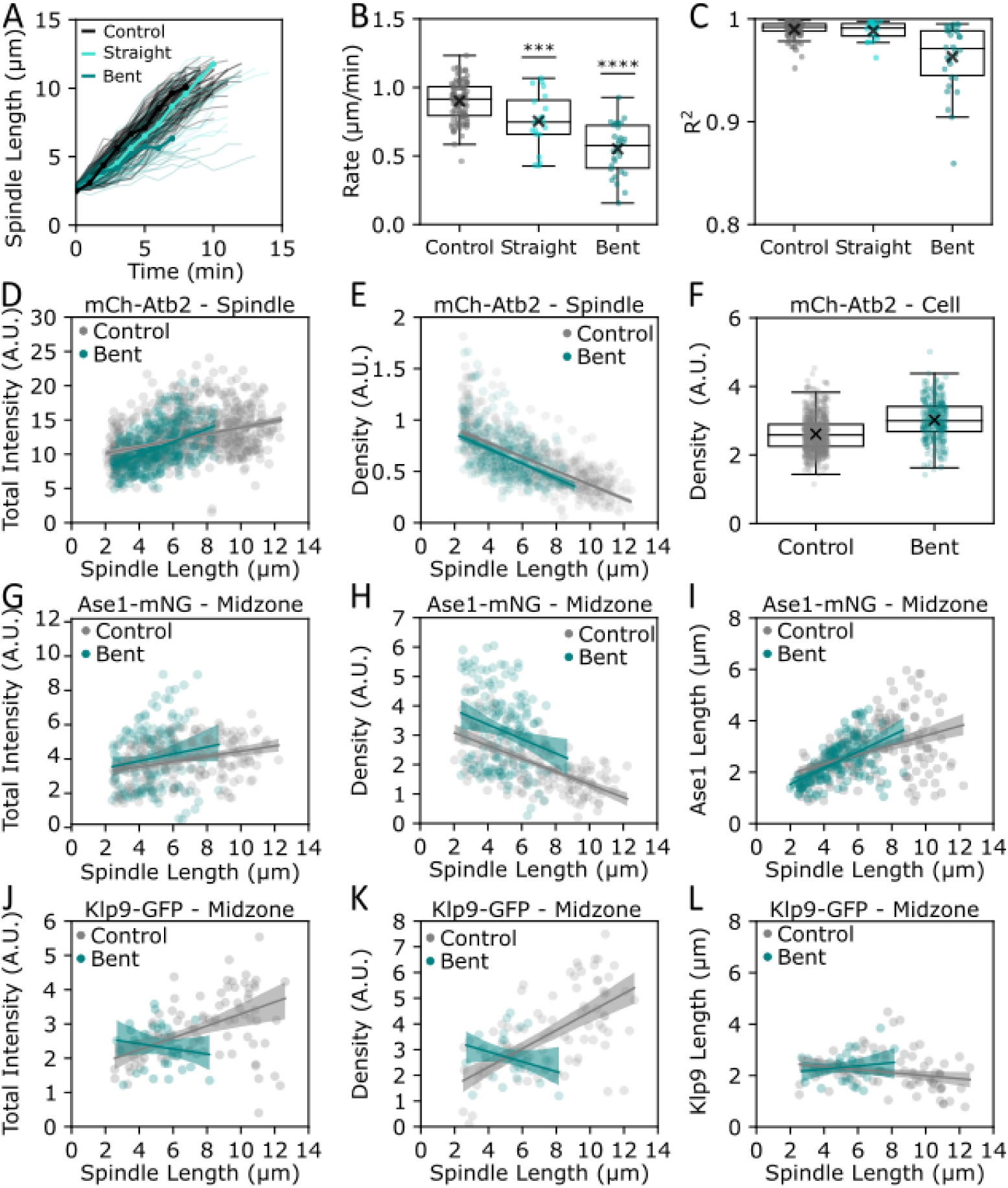
Effects of osmotic treatment on anaphase rates and quantities of spindle proteins. **(A)** Spindle elongation plots for WT cells in control conditions (black) and in cells after osmotic treatment that develop straight (light blue) or bent spindles (green). (n = 100/20/30). **(B, C)** Plot and coefficient determination (R^2^) of the mean elongation rate in control and bent spindles from (A) during anaphase B, assuming a constant elongation rate. **(D)** Summed mCh-Atb2 intensities (A.U.) over the whole spindle plotted per spindle length (µm) in control (gray) and bent (green) spindles (n = 69/52). Same data set as shown in Figure 3F, except over a full range of spindle lengths. **(E)** Linear density of mCh-Atb2 (summed intensities divided by spindle length), plotted as a function of spindle length. **(F)** Linear density of mCh-Atb2 across whole interphase cells, comparing control cells with straight spindles (gray) to treated cells with bent spindles (green). **(G)** Summed Ase1-mNG intensities in the midzone plotted as a function of spindle length. (n = 27/31). **(H)** Linear density of Ase1-mNG in the midzone (summed intensities divided by midzone length) plotted as a function of spindle length. **(I)** The size of the midzone, as depicted by Ase1-mNeonGreen localization, plotted as a function of spindle length. **(J, K, L)** Same as in (G, H, I) but with Klp9-GFP. (n = 47/23). Data in (D-L) were fitted using a linear regression model (line), shaded area CI = 95%.

WT spindles elongated from ∼2.5 µm to ∼11 µm in length over 10-12 min at a rate of 0.85 ± 0.15 µm/min (Figure 3A and 3B), consistent with previous measurements (Nabeshima et al., 1998; Krüger et al., 2021). In contrast, the bent spindles elongated from ∼2.5 µm to an average length of 6.5 µm at a significantly slower rate of 0.55 ± 0.18 µm/min, 65% of the rate of control spindles (Figure 3A and 3B). The elongation plots of individual bent spindles could be well-fitted with a linear function, indicating constant elongation rates throughout anaphase B (R² > 0.85 in all cases, Figure 3C). We also noted that in the small subset of WT cells treated with osmotic oscillations that did not buckle, anaphase rates were slower by 12% (0.75 µm ± 0.20 µm/min; Supplemental Figure S3A, S3B, and S3C), suggestive of an intermediate defect. Thus, slower anaphase rates were a feature associated with the treatment of osmotic oscillations and blue light, regardless of whether they subsequently underwent buckling. As reduced rates were observed even at the onset of anaphase B, well before spindle buckling, these data suggested that spindle bending per se was not the cause of the slowdown in the elongation rate.

One potential reason for the decrease in spindle elongation rates could be a change in the number of MTs and/or of spindle-associated proteins present in the spindle. We tested this possibility by imaging and quantifying the fluorescence intensities of mCh-Atb2 (α-tubulin), Ase1-mNG, and Klp9-GFP, which were expressed at endogenous levels from their chromosomal loci (see Methods; Figure 3D and 3E). Total mCh-Atb2 intensity in the whole spindle showed that control and bent spindles had similar numbers of MTs across a range of spindle lengths (Figure 3F and Supplemental Figure S3D and S3E). Likewise, the cellular concentrations of Atb2 tubulin were similar (Supplemental Figure S3F). In addition, the total intensities of Ase1-mNG and Klp9-GFP, as well as their localization on the midzone, were not significantly different from the control, although the values were more heterogeneous (Figure 3G and 3H and Supplemental Figure S3G-L). Thus, these data demonstrated that spindle bending and reduced anaphase rates cannot be explained simply by the numbers of MTs, Ase1, or Klp9. They further imply that the bent spindles are quantitatively similar in their architecture to the normal spindle.

### Motor mutants exhibit proportionally slower elongation rates in bent spindles

We next addressed how anaphase B elongation rates in the bent spindles were altered in representative spindle mutants. Control WT*, klp9Δ, cut7Δpkl1Δ, klp2Δ,* and *ase1Δ* mutants exhibited characteristic anaphase rates, similar to those described previously (Figure 4A and 4B) (Nabeshima et al., 1998; Loiodice et al., 2005; Bratman & Chang, 2007; Yukawa et al., 2018; Shirasugi & Sato, 2019; Yukawa et al., 2019; Loncar et al., 2020; Krüger et al., 2021). Spindle elongation was reduced by ∼50% in control *klp9Δ* and *cut7Δpkl1Δ* mutants, whereas *klp2Δ* and *ase1Δ* mutants showed elongation rates resembling those of WT.

**Figure 4.**
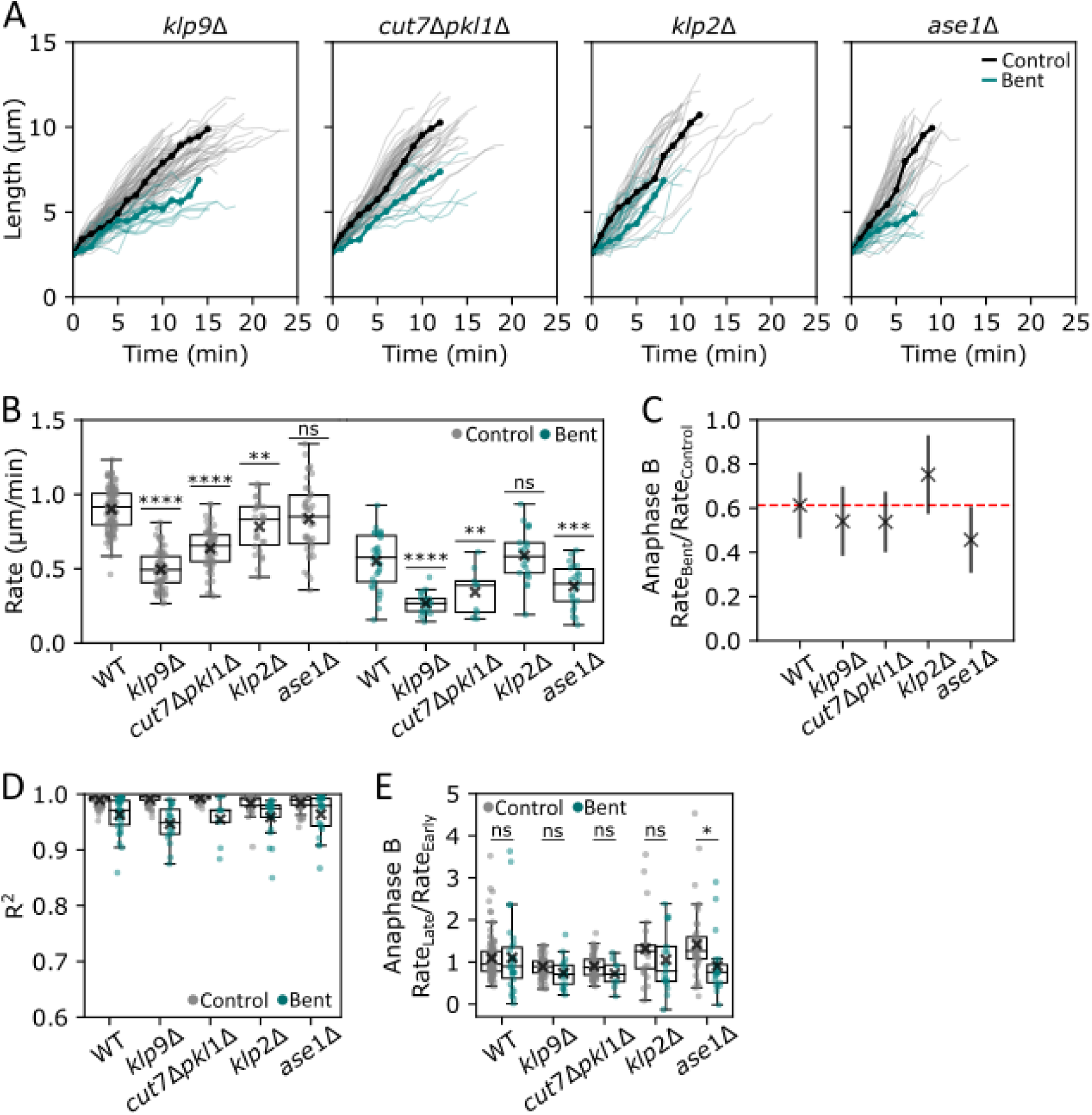
Bent spindles in mutants exhibit proportionally reduced spindle elongation rates compared to controls. **(A)** Spindle elongation plots in control (grey) and bent spindles (green) for the indicated spindle mutants. Each line shows one spindle; representative spindles are highlighted in bold (n = 71 /56/26/41; 19/9/21/22). **(B)** Mean spindle elongation rates from spindles in (A). **(C)** Ratio of bent to control spindle elongation rates (µm/min). The red line represents the predicted ratio according to a power model (see Methods). **(D)** Coefficient of determination (R^2^) for all the individual plots in (A) fitted to a linear regression model assuming constant elongation. **(E)** Ratio of elongation rates in early versus late anaphase B.

The bent spindles in these strains all exhibited slower anaphase rates than their untreated straight counterparts (Figure 4A and 4B). Interestingly, the reduction in rates for control vs. bent spindles was proportional: the bent spindle rates were approximately 60% (range 55-74%) of the controls for WT, *klp9Δ,* and *cut7Δpkl1Δ* mutant strains (Figure 4C). Notably, the bent *ase1Δ* mutant spindles showed a stronger rate reduction than the other mutants, which may reflect its compromised structural stability.

Across these strains, elongation rates were largely constant throughout anaphase in both bent and control straight spindles. The plots of spindle length as a function of time fit well to a linear model (Figure 4D). In addition, by comparing the elongation rates in early and late anaphase B, we did not detect anaphase stalling up to the point of spindle breakage (Figure 4E). The slower elongation rates support a model in which anaphase spindles in cells treated with osmotic oscillations and blue light work against an increase in mechanical load that remains constant throughout anaphase B (see Methods).

### Spindles break at a consistent size, not time

We sought to understand the factors determining when the spindle breaks or disassembles. We considered two models: spindles may break or disassemble at a specific size (“sizer”) or after a specific period of time (“timer”); these are conceptually analogous to “sizer” vs “timer” models in cell size homeostasis (Facchetti et al., 2017; Miller et al., 2023).

To test these models, we measured the size of WT and mutant spindles that elongated at different rates. Control straight WT spindles all elongated to a terminal size of 9-11 µm before disassembly, with subtle size differences in the mutants (Figure 5A). Bent spindles broke at 6-7 µm in WT and mutants, except for *ase1Δ* mutants that broke below 5 µm (Figure 5B). In contrast, the time in anaphase until breakage or disassembly varied significantly across the strains (Figures 5C and 5D). Furthermore, compiling the spindle lifetime against the range of elongation rates in WT and mutant cells in a single graph showed that the time until spindle breakage or disassembly negatively correlates with the anaphase elongation rate (Figure 5E). These results indicate that spindles break or disassemble at a specific spindle size and not time, supporting a sizer mechanism rather than a timer mechanism. One explanation for this size-dependent effect could be the accumulation of mechanical load in the spring-like spindle that triggers breakage once a critical size and corresponding critical load are reached.

**Figure 5.**
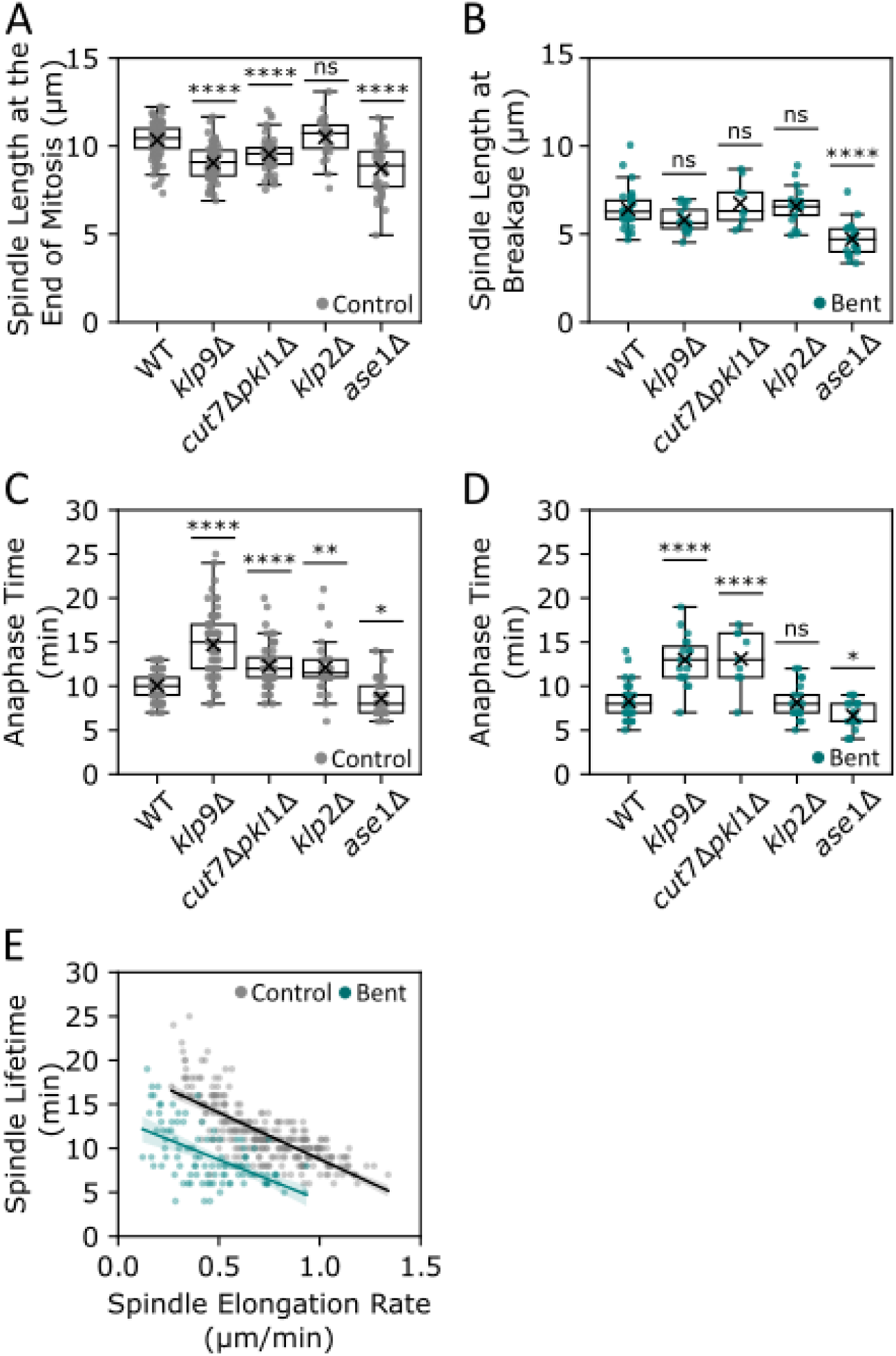
Spindles break or disassemble at a constant size, not time. **(A)** Spindle length (µm) just before spindle disassembly at the end of mitosis. (n = 100/71/56/26 41) **(B)** Spindle length just before spindle breakage in bent spindles. (n = 30/19/9/21/22). **(C)** Total time (min) in anaphase B in control spindles from (A). **(D)** Time in anaphase B to spindle breakage in bent spindles from (B). **(E)** Inverse relationship of anaphase spindle lifetime with elongation rate, using data from WT and mutant cells from (A-D). Each point represents a cell. Linear fit (line) with shaded area CI = 95%.

### The spindle behaves as a beam with increased rigidity in the midzone

To analyze the material properties of the fission yeast spindle, we measured and analyzed the shapes of the bent spindles. The spindle may be a mechanically more complex object than a homogeneous beam because of the heterogeneous organization of MTs and spindle factors along its length. For example, the midzone region may be more rigid than the rest of the spindle in part due to the higher number of overlapping MTs. Electron tomography images of mid-anaphase spindles ∼5 μm in length show that the midzone region consists of 6-9 anti-parallel overlapping MTs organized in a square-packed array, while the surrounding region consists of parallel bundles of 3-5 MTs organized in hexagonal packing (Ding et al., 1993; Ward et al., 2014). Modeling predicts these differences in MT number and packing within the mid-anaphase spindle may lead to anywhere from a 2 to 10-fold increase in rigidity at the midzone, depending on the parameters used (Ward et al., 2014). In addition, rigidity may be further modulated by the binding of spindle factors such as Ase1 (PRC1/MAP65), which may potentially alter the local rigidity of individual MTs as well as MT bundles (Portran et al., 2013).

To study the bending rigidity of spindles *in vivo*, we extracted the shape of each bent spindle and the extent of the midzone region from images of spindles marked by mCh-Atb2 and Ase1-mNG (Figure 6A, Supplemental Figure S6B; see Methods). We developed a model to simulate the shape of a bent beam with the same pole-to-pole distance and spindle length, allowing the rigidity in the midzone region to vary relative to the rest of the spindle (see Methods; Figure 6A). Comparison of experimental and theoretical shapes allowed us to assess the difference in rigidity between the midzone and the rest of the spindle as a rigidity ratio “r”.

**Figure 6.**
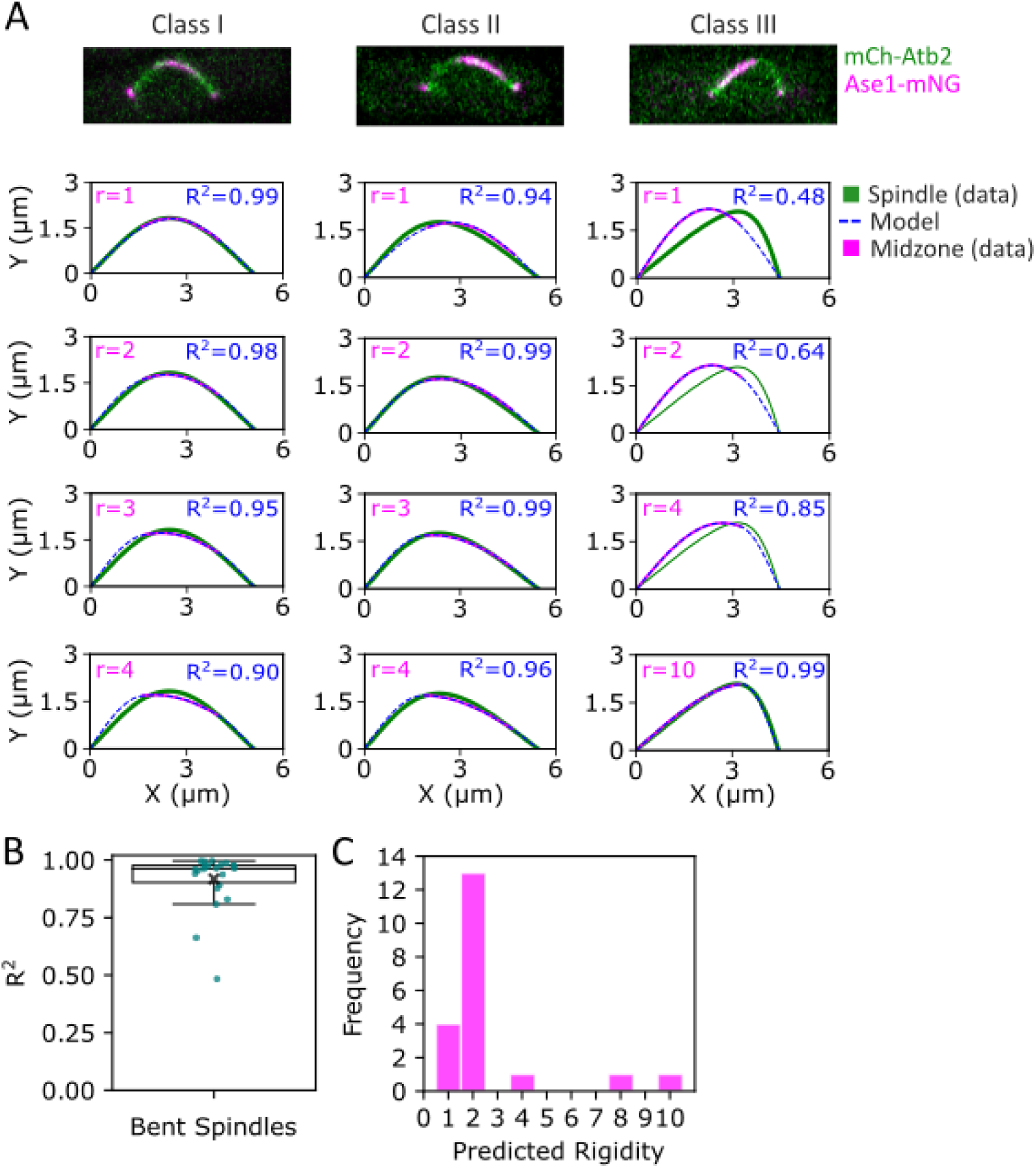
The spindle behaves as a non-homogeneous beam with increased rigidity in the midzone. **(A)** confocal maximum intensity projection images of bent spindles tagged with mCh-Atb2 and Ase1-mNG, representative of three Classes. The shape of each experimental spindle (green) was compared with modeled spindles (dashed blue line) with the midzone region (magenta) to yield a coefficient of determination (R^2^) for each comparison. In the model, the rigidity ratio (r) of the midzone/non-midzone regions of the spindle was varied to determine what rigidity ratio value produced the best fit for each spindle. **(B)** Coefficient of determination (R^2^) comparing experimental spindle shapes with the modeled homogenous spindle shapes where r = 1 (n = 22). **(C)** Predicted midzone rigidity ratios based upon the shape of bent spindles (n = 20).

**Supplemental Figure S6.**
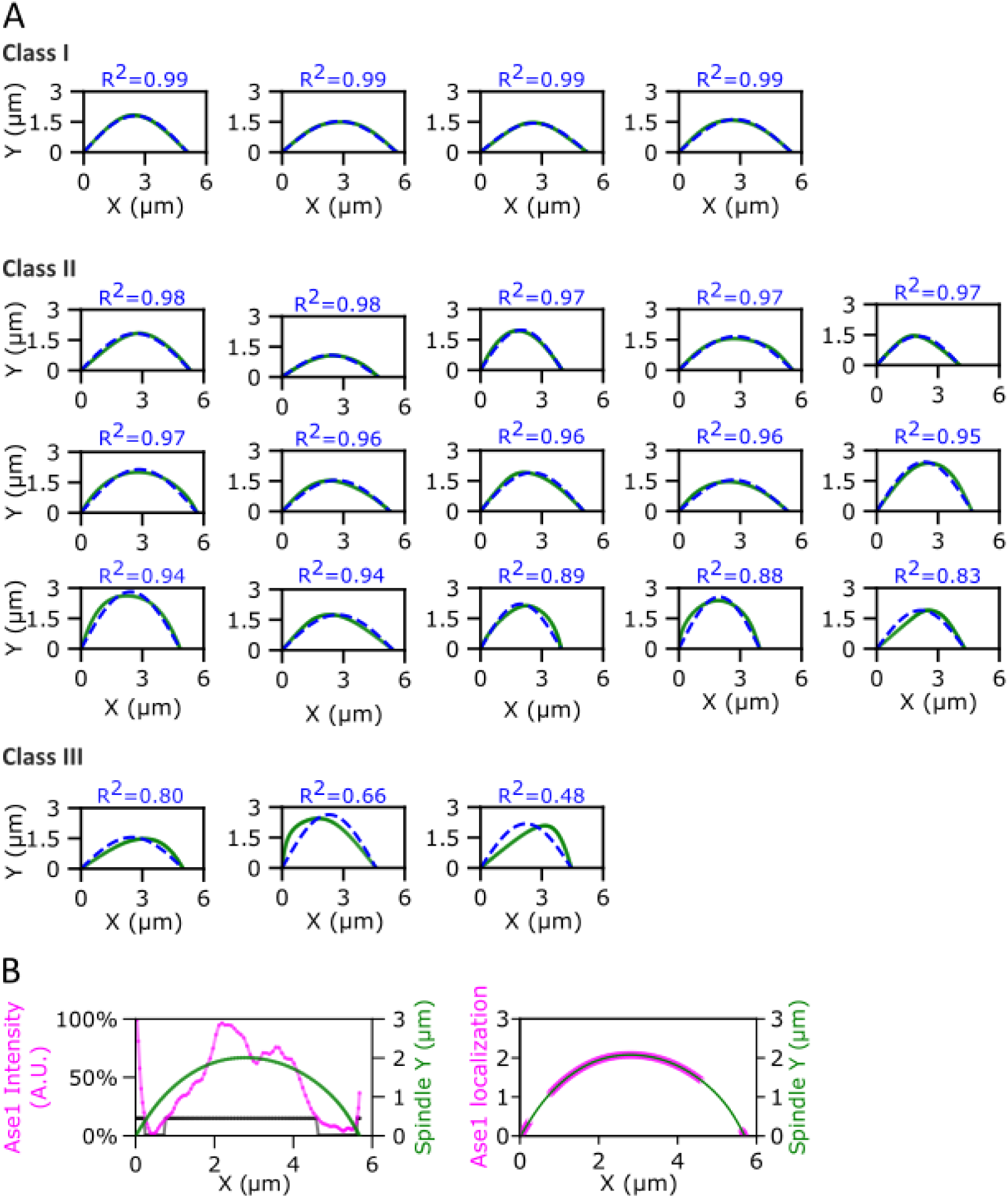
Comparison of the shapes of experimental bent spindles with the model of a homogeneous beam. **(A)** Graphical representations of 22 experimental bent spindles (green) compared to a modeled homogeneous beam (dashed blue) with the same length and pole-to-pole distance as the experimental spindle. Spindles are grouped by class and ordered (left to right and top to bottom) by decreasing coefficient of determination (R^2^) comparing the experimental and modeled spindle shapes. **(B)** Example of how Ase1 (Ase1-mNG) localization was input into the model. The left graph shows Ase1-mNG intensity profile (magenta) along the spindle length (green) and the 15% of intensity threshold applied (black) to determine its localization. The right graph shows how the Ase1 localization (magenta) would look on the corresponding spindle (green) after applying the threshold.

We grouped the spindles into three distinct classes based upon how closely they fit the shape of a homogenous beam (r = 1) (Figure 6A, 6B, 6C, and Supplemental Figure S6A). 20% of the spindles (n = 20) closely approximated the theoretical shape of a bent beam with homogeneous rigidity throughout its length (Class I; R^2^ > 99% when r = 1). In this category, the fit between the simulation and experiment was not clearly improved by increasing the rigidity of the midzone. 65% of the spindles exhibited a mild deviation from the homogeneous beam model (Class II; R^2^ > 99% when r = 2). In these spindles, the fit was improved to a model with a two-fold increase in rigidity in the midzone region. Most of these spindles were slightly asymmetric: they appeared slightly tilted to one pole and exhibited a midzone region that was longer, flatter, and shifted towards the opposite pole. Finally, 15% of the spindles compared poorly to the homogeneous beam (Class III; R^2^ > 99% when r ≥ 4). These shapes fit well with models with a large increase in midzone rigidity; alternatively, the shape of these spindles may also be explained by additional forces on the spindle, such as from chromosomes or the NE. These bent spindles were characterized by large, asymmetric flattened midzone regions. In summary, these data indicate that a large majority of the bent spindles (85%, Class I and II) behaved as near-homogenous beams with only a mild 2-fold or less increase in rigidity in the midzone region (Figure 6D).

### Spindles break at a fragile site located near the edge of the mid-zone

Finally, we examined the mechanism by which bent spindles break. We first hypothesized that the spindle would break at or near the site of greatest local curvature, corresponding to the region of maximal bending strain. To map sites of breakage, we imaged bent spindles labeled with mCh-Atb2 and Ase1-mNG at high frequency (every 6 sec) (Figure 7A). The bent spindles broke across a wide range of maximal local curvatures (Figures 7B and 7C), suggesting that there was no consistent curvature threshold. Furthermore, spindles did not break at their site of maximal local curvature, but instead at some distance away (Figures 7B and Supplemental Figure S7A).

**Figure 7.**
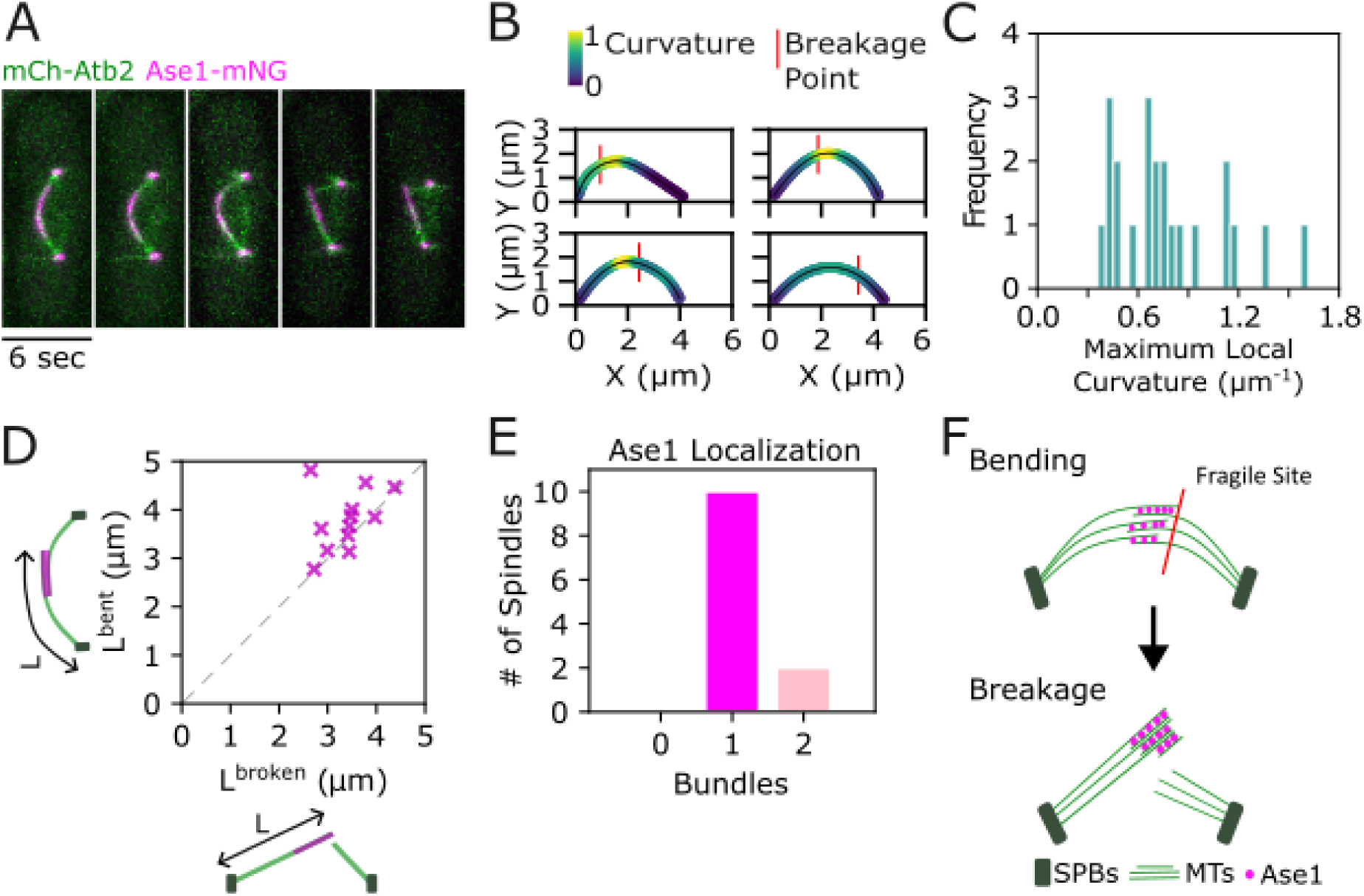
Spindles break at a fragile site located near the edge of the midzone. **(A)** Time-lapse maximum intensity projection images of a bent spindle breaking, with MTs (mCh-Atb2, green) and Ase1 (Ase1-mNG, magenta) labeled. See Supplemental Figure S7 for additional examples. **(B)** Comparison of bent spindle curvature (normalized) and the subsequent breakage point (red) in 4 representative spindles. **(C)** Maximum curvature (µm^−1^) of bent spindles at the time point just before spindle breakage (n = 22). **(D)** Bundle length (µm) from the SPB to the edge of the midzone before (L^bent^) and after breakage (L^broken^) (t = 6). Data were fitted to a linear regression model (line) with CI = 95% (shaded areas) (n = 11). **(E)** Localization of Ase1 just after spindle breakage to spindle bundle remnants. (n = 12). **(F)** Proposed model for the breakage of bent spindles at the edge of the midzone.

**Supplemental Figure S7.**
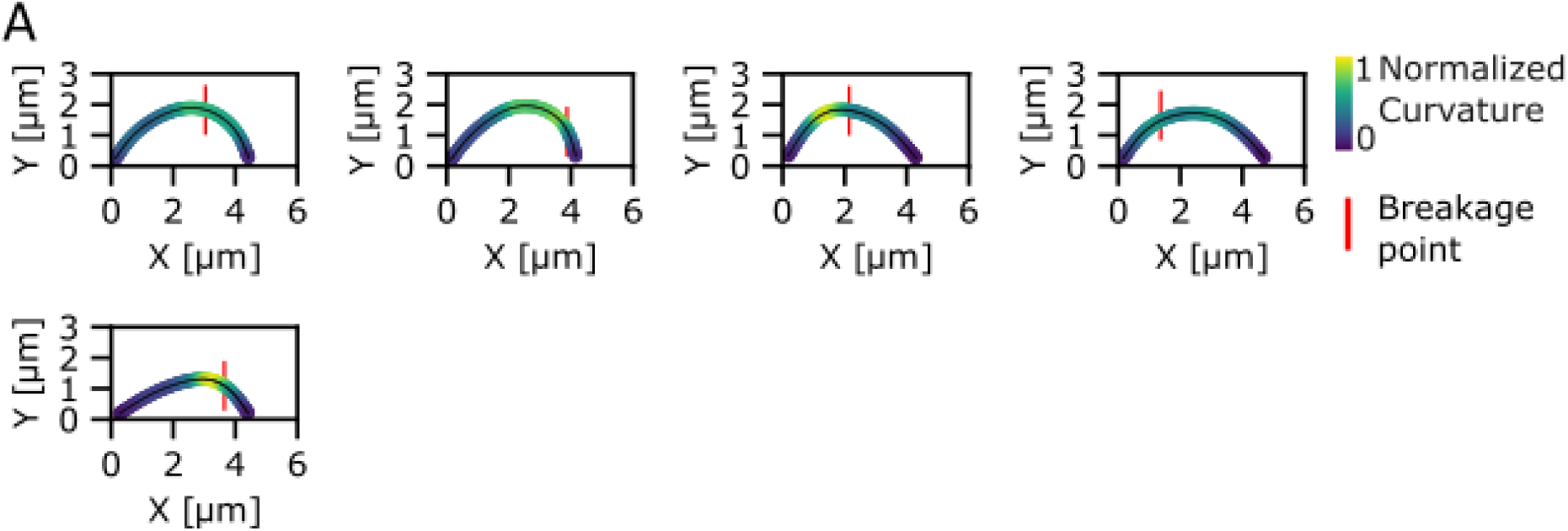
Bent spindles do not break at the point of maximal local curvature. **(A)** Comparison of bent spindle curvature (normalized) and the subsequent breakage point (red). These are additional examples to those shown in Figure 7B.

Closer analysis revealed that nearly all spindles (89%) broke near the edge of the midzone, which was defined by Ase1-mNG localization (Figure 7D). In most cases (83%), the spindle broke at the end of the spindle midzone, producing two different-sized bundles (Figure 7D). Typically, the larger spindle fragment retained the Ase1-mNG-decorated region with no diminution of the Ase1-mNG intensity; this fragment grew rapidly with Ase1 maintained at the end, often extending past the SPB. The shorter spindle fragment generally did not contain detectable Ase1-mNG and quickly shrank. In 11% of cases, the spindle broke within the midzone, so that Ase1 was localized on both fragments, which were initially stabilized (Figure 7E). In general, these behaviors were similar to those of spindle fragments generated in other ways (Khodjakov et al., 2004; Zheng et al., 2007).

Together, our observations suggest a model for spindle breakage: mechanical strain causes the simultaneous, catastrophic fracture of multiple MTs at a fragile site on the edge of the midzone (Figure 7F). One of the spindle fragments retains the midzone proteins, such as Ase1, and likely other MT stabilizing proteins, such as Klp9 and the CLASP orthologue (Bratman & Chang, 2007; Ebina et al., 2019; Yukawa, Okazaki, et al., 2019; Krüger et al., 2021), which allows the fragment to elongate. In contrast, the remaining MT fragment, lacking stabilizing proteins, rapidly depolymerizes and shrinks. These findings indicate that these MTs at the edge of the midzone represent the weakest point in the bent spindle, more fragile than the MT crosslinkers, NE, or chromosomes.

## Discussion

### An approach to interrogate spindle mechanics

In this study, we investigated the buckling behavior of the *S. pombe* spindle to understand how spindles elongate, bend, and break under mechanical load. Our experiments revealed that treatment with osmotic oscillations and blue light induced anaphase failure (Figure 1A and 1B), highlighting how these environmental perturbations can produce significant and specific effects on mitosis. We used this protocol to generate and analyze large numbers of bent spindles, as it consistently produced more penetrant and pronounced phenotypes than those generated by other methods (Nakazawa et al., 2016; Begley et al., 2025). For example, the drug cerulenin, an inhibitor of fatty acid synthase, results in spindles that buckle to varying extents, with only a subset actually breaking (Yam et al., 2011). In contrast, our approach produced bent spindles in over 75% of mitotic cells, with all these spindles bending and ultimately breaking, resulting in mitotic failure. Our quantitative studies were thus facilitated by the high penetrance and reproducibility of this phenotype.

In these studies, we elaborate upon a mechanical-based model in which spindle buckling and breaking are driven by spindle elongation forces opposed by resistive forces. The factors driving spindle elongation include the motor proteins Klp9 (kinesin-6), which may contribute to both MT sliding and polymerization at the midzone, and Cut7 (kinesin-5) (Hagan & Yanagida, 1992; Yukawa et al., 2018; Yukawa, Okazaki, et al., 2019; Krüger et al., 2021). Loss of either factor slowed spindle elongation but did not abolish buckling, indicating the spindle still generates sufficient force to bend itself. On the other hand, Ase1 (PRC1 orthologue) was required for the structural integrity of the spindle under mechanical load, as in *ase1Δ* mutants, the bent spindle collapsed with increased load. On the molecular level, Ase1 likely contributes multiple roles: a MT bundler, a brake resisting MT sliding (Loiodice et al., 2005; Lansky et al., 2015), and a scaffold that organizes midzone proteins, such as Klp9 and the CLASP Cls1/Peg1 (Bratman & Chang, 2007; Fu et al., 2009b; Ebina et al., 2019; Jagrić et al., 2021; Lera-Ramirez et al., 2022).

### Beam buckling mechanics

An original rationale for studying spindle buckling is to use the principles of beam buckling mechanics to estimate the forces responsible for anaphase B spindle elongation. Indeed, our findings show that the fission yeast spindle can be regarded as a beam that buckles while anchored only at both poles. The analyses of spindle shapes showed that a subset of the bent spindles closely approximates the shape of a buckled homogenous beam (Figure 6). However, most of the spindles fit optimally to a model of a non-homogeneous beam with a modest (two-fold or less) increase in rigidity at the midzone (Figure 6). This result contrasts with some models suggesting that the midzone may have up to a ten-fold increase in rigidity due to differences in MT number and structural packing (Ward et al., 2014).

Using parameters such as beam dimensions and material properties, simple beam mechanics can estimate the force required to bend the spindle. In their initial models, Ward et al. (2014) suggest that the critical buckling force relates to spindle length of F_c_ ∼ L^−4^ and provide an initial estimate of 31 pN spindle elongation force needed to buckle a fission yeast spindle ∼5 µm in length (Ward et al., 2014). However, there are critical parameters that remain unquantified in a rigorous manner, notably the precise number of MTs present in the spindle. Fluorescence intensity measurements indicate that the numbers of MTs, as well as the proteins Ase1 and Klp9, in bent spindles are similar to those in control spindles (Figure 3). Unfortunately, measurements of MT numbers in wild-type control spindles have been inconsistent across different studies (Ward et al., 2014; Loiodice et al., 2019) (see Methods). This inconsistency underscores the need for further research to establish quantitative estimates of the forces at play during anaphase B.

### Forces resistive to spindle elongation

In addition to forces that elongate the spindle, our study also highlights the existence of forces that resist spindle elongation in the bent spindle. These forces, which are increased by exposure to osmotic oscillations and blue light during interphase, not only limit pole-to-pole separation at mid-anaphase but also slow down the rate of spindle elongation throughout anaphase B. The presence of intermediate phenotypes suggests that these resistive forces may be tunable (Supplemental Figure S1).

Although the precise origins of these resistive forces remain unknown, two leading candidates merit discussion: forces related to chromosome segregation and those associated with nuclear envelope expansion. It is well established that active forces are necessary for the proper separation of sister chromatids during anaphase (Jannink et al., 1996; Lawrimore & Bloom, 2022). Previous studies have demonstrated that peri-centromeric DNA is organized into loops into a bottlebrush structure by cohesins and condensins, which must be actively disassembled and untangled by topoisomerase 2 during anaphase to achieve successful chromosome segregation (Lawrimore & Bloom, 2022; Kolbin et al., 2025). Chromosomal regions undergo stretching during anaphase caused by residual cohesion between sister chromatids, followed by a recoil, indicating the presence of chromosome-based resistive forces even in normal cells (Renshaw et al., 2010).

Our various observations are consistent with defects in chromosome separation, including the presence of anaphase DNA bridges, a decrease in anaphase elongation rates, and bent spindles (Figures 1 and 2), which are also seen upon topoisomerase 2 inhibition (Okamoto et al., 2012; Nakazawa et al., 2016). The effects of blue light as a source of genotoxic stress are well documented (Chamayou-Robert et al., 2022; Camponeschi et al., 2023). Osmotic oscillations, which induce repeated fluctuations in nuclear dimensions, may disrupt DNA movement and topology. Therefore, we speculate that the combination of blue light exposure and osmotic oscillations creates synergistic effects on increased DNA entanglement during S phase and DNA repair, which cause an increase in chromosome-based resistive forces and ultimately lead to mid-anaphase failure.

Another potential source of resistive forces during cell division is NE tension. Previous studies suggest that cerulenin treatment and mutations affecting lipid metabolism contribute to spindle buckling in the closed mitosis of *S. pombe* by inhibiting the NE’s ability to expand properly during mitosis (Yam et al., 2011; Takemoto et al., 2016; Expósito-Serrano et al., 2020; Lera-Ramirez et al., 2022; Foo et al., 2023; Begley et al., 2025). Support for this view comes from experiments demonstrating that the natural breakage of the NE in *S. japonicus* (Yam et al., 2011) and laser cutting near the NE in cerulenin-treated *S. pombe* (Begley et al., 2025) release bent spindles into a straight configuration, facilitating successful division. However, it remains unclear whether these spindle buckling effects arise from a physical limitation in the available nuclear membrane or from some other underlying defect. Notably, *S. pombe* cells possess significant stores of membrane available for NE expansion, potentially derived from the endoplasmic reticulum (ER), as they can acutely expand their nuclear envelope by 40% in surface area when exposed to hypoosmotic shock (Lemière et al., 2022). Furthermore, cerulenin treatment does not inhibit the normal growth of the nucleus during interphase (Yam et al., 2011; Kume et al., 2019). As an alternative explanation, we suggest that the effects of cerulenin and lipid metabolism on spindle buckling may also be explained by emerging evidence highlighting the role of lipids and the nuclear envelope in regulating chromatin organization (Sosa Ponce et al., 2025).

Overall, our observations currently favor a chromosomal separation defect as the underlying cause of these phenotypes. Although NE restriction cannot be ruled out, it does not readily explain why effects are only manifest during mitosis, or why spindle elongation rates are decreased in early anaphase. As we focused primarily on spindle mechanics in this study, future experiments are needed to define the specific mechanisms causing this intriguing phenotype.

### Identification of a spindle fragile site

Spindles can encounter mechanical perturbations from their environment, as well as from chromosome segregation defects that can end in spindle breakage (Valdez et al., 2023). Our studies showed that the bent spindles broke in a highly stereotypical manner, indicative of a fragile spindle site. Mechanical failure occurred at the edge of the spindle midzone, at which multiple MTs in a parallel bundle appear to fracture near-simultaneously (Figure 7). The preference for this site suggests that it is more fragile than other parts of the spindle, such as the MT crosslinkers, chromosomes, or the nuclear envelope. This fragile site likely represents a transition of a parallel bundle of 3-4 MTs to an anti-parallel bundle of 6-8 MTs bound by midzone proteins (Ward et al., 2014). This fragility at a point of mechanical discontinuity may be analogous to the mechanism by which F-actin is severed by cofilin at boundary sites between bare and decorated segments, and reflects general properties of fracture sites in materials (de la Cruz, 2009). Recent modeling of MT breakage by mechanical stress produced by clusters of kinesin proteins estimates that the rupture force for an individual MT is on the order of 70-150 pN (Geng et al., 2025). Thus, although it is unknown how discontinuity influences the force requirement, we predict that quite large forces are required to break a bundle of MTs.

The breakage of the bent spindles occurred at a consistent spindle size independent of elongation rate, suggesting a “sizer” rather than a “timer” mechanism (Figure 5). One explanation for this size dependence could be that the accumulation of mechanical strain reaches a critical point at these specific dimensions of the bent spindle (see Methods). Although no MT severing proteins have been identified in fission yeast, it is possible that other regulatory mechanisms at the end of anaphase (Salas-Pino & Daga, 2019) may also sense mechanical or size cues to stimulate the disassembly of these bent spindles.

### Relevance to large complex animal spindles

One challenge in the study of mitosis in animal cells is the sheer complexity of the spindle, which is composed of 10,000 to 100,000 MTs (Nicklas, 1983; Brugués & Needleman, 2014; Yu et al., 2019; O’Toole et al., 2020; Conway et al., 2022; Kiewisz et al., 2022). In animal cell models, multiple parts of the spindle contribute to anaphase B forces, including pushing forces from bridging fibers connected to kinetochore fibers within the spindle, k-fiber MT depolymerization at the poles, and pulling forces from astral MTs on spindle poles (Vukušić et al., 2019; Anjur-Dietrich et al., 2021; Valdez et al., 2023). Other forces impacting chromosome segregation include forces from the cytoplasm and cytoplasmic flows (Anjur-Dietrich et al., 2021; Xie et al., 2022, 2025) as well as potentially interactions with organelles such as the ER, NE, and condensates (Scholey, 2025). The minimal yeast spindle is likely to be analogous to bridging spindle fibers that contribute to pushing forces for anaphase B in animal cells (Tolić, 2018), and thus these studies hold direct relevance towards understanding a key portion of complex animal spindles.

## Material and Methods

### Yeast strains and growth conditions

All *S. pombe* strains are listed in Table S1. Standard fission yeast molecular genetics techniques and media were as described (Moreno et al., 1991).

The Ase1-mNG gene fusion was constructed by targeting mNeonGreen-kanMX into the Ase1 chromosomal locus using the pFA6A mNeonGreen-KanMX plasmid (Addgene plasmid # 129099; http://n2t.net/addgene:129099; RRID: Addgene_129099). The PCR primers were: 5’-CTACCAACATTTTTTCTGCTCCACTCAACAATATTACAAATTGTACACCGATGGAGGATGAATGGGGAGAAGAAGGCTTTATGGTGAGCAAGGGCGAGGA-3’ and 5’-TGAATGCTGGTCGCTATACTGGCTTCTTATTTACCTAATCGATCAAATTTAAATATACATATTTTTGCATATGAATACAGCATATAGATAATTCATAAAA-3’.

### Microscopy

Cells were imaged on a customized microscope system that includes a Ti-Eclipse inverted stand (Nikon Instruments) with perfect focus, a spinning-disk confocal head (Yokogawa, CSU-10), 488 nm and 541 nm laser lines illuminated controlled by Borealis system, with emission filters 525 ±25 nm and 600 ±25 nm respectively, a 60X (NA: 1.4) Plan Fluor objective with 2.5x magnifier, EM-CCD camera (Hamamatsu, C9100-13) and a piezo-controlled XYZ stage. These components were controlled with µManager v. 1.41 (Edelstein et al., 2010). A black panel cage incubation system (OkoLab, #748-3040) was used to control temperature.

### Osmotic oscillation protocol to generate bent spindles

Cells were grown in CellASIC microfluidic chambers (designed for 3.5-5 µm haploid yeast cells, Millipore Sigma, Y04C-02-5PK) controlled by the ONIX2 microfluidic system (Millipore Sigma, CAX2-S000) for 5 min. These chambers were first prewashed with YES media at 8 psi.

Fission yeast cells were grown overnight in liquid cultures at 30°C in YES medium to the exponential phase and then introduced into the CellASIC chambers at 30°C with a continuous flow of media at a 5 psi setting. Cells were first grown in YES medium for a 20-min adaptation period. Then, cells were treated to 5 osmotic oscillatory cycles, where each cycle consisted of 5 min of YES followed by 5 min of YES + 1 M sorbitol. After these osmotic oscillations, cells were grown in YES for the rest of the experiment. During the first 180 min (including the oscillations), images in 20-30 fields of view were acquired every 10 min. Following this, 4-5 fields of view were imaged every min for 30 min. This last step was repeated 4-5 times over 2-2.5 hr to image multiple other fields of view, each for 30 min, to capture additional mitotic events. The sequential sampling of different fields allowed us to image mitotic events from roughly 20 fields of view at a range of time points after the oscillations.

During this protocol, cells were imaged using a spinning disc confocal microscope and a brightfield microscope to track cell cycle progression. At each time point, a stack of 13 Z planes 0.5 μm apart was acquired, one for each laser wavelength. Cells were exposed to a 488 nm laser (50 mW at 2% power) with 50 ms acquisition per Z plane, and to a 561 nm laser (50 mW at 5% power) with 50 ms acquisition per Z plane. We estimated that cells were exposed to > 600 acquisitions for each laser over the course of each experiment.

For spindle breakage analysis, Z-stacks were acquired every 6 s to capture the rapid dynamics over 120 s (Figures 2B, 2C-*ase1Δ*, and 7).

### Measurement of spindle dimensions and elongation rate

Confocal time-lapse images of spindles were analyzed using Fiji ImageJ (Schindelin et al., 2012). Spindles were manually curated based on the following criteria: (i) all portions of the spindle remained within the central focal plane (with 3 central Z planes of 0.5 µm) during the time period of analysis; (ii) the entire anaphase B period was captured, starting from anaphase B onset at spindle length of 2.5 ±0.1 µm (Kruger et al., 2019) to spindle disassembly or breakage. A maximum intensity projection of the Z-stack images of mCh-Atb2 or GFP-Atb2 was used for the determination of spindle length. In contrast, images of Mis12-GFP foci were used to determine the pole-to-pole distance. Elongation rates were derived from spindle lengths taken from time-lapse images 1 min apart. Spindle lengths were measured in Fiji by manually drawing a straight or curved line over each spindle for each time point. The SciPy Python package (Virtanen et al., 2020) was used to fit the length (L) versus time (t) for each individual spindle to a linear regression and to calculate each coefficient of determination (R^2^) and elongation rate (slope dL/dt) for each spindle (Figure 3, Figure 4, and Figure 5E)

### Photobleaching and fluorescence intensity measurements

The fluorescence intensities of mCh-Atb2, Klp9-GFP, and Ase1-mNG were measured on SUM intensity projections of time-lapse spindle images curated as described above (Figure 3). For mCh-Atb2 and Ase1-mNG, Z-stacks were acquired every 1 min for 30 min, and for Klp9-GFP, every 5 min for 30 min. Tubulin (mCh-Atb2) intensity was measured within a 0.45 µm-wide line region of interest over the whole spindle, while Klp9 and Ase1 intensities were measured within a 0.45 µm-wide line region of interest at the midzone only. “Total Intensity” was calculated by summing the intensities of the pixels in the ROI, and “Density” was calculated by dividing the sum of the pixels in the ROI by the ROI area. These intensity values were corrected by background subtraction and for photobleaching. For background subtraction, the intensity of a region of interest inside the nucleus that did not include the spindle was measured for each spindle. To quantify photobleaching, the intensities of each fluorescent protein were measured over multiple fields of view. Photobleaching was quantified by fitting a single exponential decay of fluorescence signal using the SciPy Python package and a custom Python script. Calibrations showed that fluorescence intensities remained consistent across different days.

### Analyses of spindle shape

For Figures 6 and 7, spindles were manually traced along their entire length using mCh-Atb2 SUM projection images. Using the segmented line tool and spline fit in Fiji, we recorded 100 equally spaced coordinates along each spindle. A custom Python script was then used to translate and rotate the spindle coordinates so that all spindles were aligned with their poles on the horizontal axis, with one pole positioned at (0,0) and the second at a distance D on the same axis (D,0). Once aligned, the traces were fit to polynomial regression models, increasing the polynomial order until the coefficient of determination (R²) between the trace and the polynomial exceeded 0.9999, at which point the fit was considered sufficiently accurate. The resulting polynomial equation was then used to measure spindle length by computing the arc length between the poles.

### Comparison of spindle shape with beam buckling theory

To compare the spindle deformation with the behavior of a beam that spontaneously bends from straight to curved under load, we use the simplest case of a homogeneous and isotropic beam buckling, governed by the Euler-Bernoulli equation. We impose boundary conditions that allow both ends of the beam to rotate freely. The model was restricted to the first buckling mode (n=1), and a fixed pole-to-pole distance (D) and the experimental spindle length for each spindle. The solutions for the buckling shape were given by:

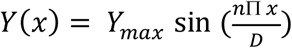

where *Y*_*max*_ represents the maximum lateral deflection.

### Modeling midzone localization

To determine the localization of the midzone for each spindle analyzed, we assess the fluorescence intensity distribution of Ase1-mNG. The manual spindle trace obtained from the mCh-Atb2 channel was also used to measure Ase1-mNG intensity along the same spindle. Using a custom Python script, we aligned the spindle coordinates with the Ase1 intensity signal. To achieve this, the Ase1 intensity was interpolated onto the 100 equally spaced coordinates along the x-axis projection of the spindle. This ensured that for each given x-coordinate, both the spindle deflection and Ase1 intensity were known. To minimize bias from the strong Ase1 signal at the spindle poles, we filtered the intensity signal by measuring the maximum intensity within the central 60% of the spindle. A threshold was then set at 15% of this maximum intensity value—regions with an intensity above this threshold were classified as Ase1-positive, while regions below were considered Ase1-negative. This filtering method was applied to localize the presence or absence of Ase1 along the spindle.

### Non-homogeneous beam theory simulation

To model the spindle as an isotropic beam with non-uniform rigidity along its length, we considered the case where the midzone, corresponding to the region of MTs overlap, exhibits increased rigidity. To compute the Euler buckling profile for such a heterogeneous beam, we used the colbuc function from Advanced Mathematics and Applications Using MATLAB (Howard Wilson, 2025), which allows for variation in rigidity along the beam length. We used the Ase1-mNG localization profile to identify the spindle midzone and defined this region as the section with higher rigidity. The model parameters included the total spindle length, the pole-to-pole distance, the extent of the midzone, and a rigidity ratio (r) between the midzone and the rest of the spindle. For each spindle, we iteratively increased the rigidity ratio (r) in steps of 1 and computed the corresponding theoretical buckling shape. We then quantified the agreement between the experimental spindle shape and the model prediction by calculating the coefficient of determination (R^2^) between the experimental trace and the theoretical profile.

### Local curvature measurements

Local curvature along the spindle was computed using a custom Python script adapted from (Suresh et al., 2020) on the manually traced spindle shapes derived from spindle images. Briefly, for each point along the spindle, we calculated the radius of a circle passing through three points spaced five pixels apart (0.55 µm). This spacing corresponds to a biologically relevant length scale (in microns) for assessing spindle bending. The local curvature was then defined as the inverse of this radius and mapped along the spindle trace. Due to the five-pixel spacing required for the calculation, curvature values could not be computed for the first and last five coordinates of each spindle trace.

### Statistical analysis

Statistical tests were performed using the SciPy Python package and an Unpaired or Welch’s t-test (Figure 1E, 3B, and 4E and Supplemental Figure S1D and S1E) or a One-Way ANOVA test followed by multiple comparisons (Figure 2C, 4B, 5A, and 5B and Supplemental Figure S3B) in GraphPad Prism version 10.2.2 for Windows (GraphPad Software, Boston, Massachusetts, USA). Stars denote the significance: ns = no significance p>0.05; * p ≤ 0.05; ** p ≤ 0.01; *** p ≤ 0.001; and **** p ≤ 0.0001.

Whisker plots display the first quartile (Q1) with the bottom line, the median (Q2) as a line within the box, and the third quartile (Q3) as the top bar of the box. The ‘X’ denotes the mean, and the whiskers represent the standard deviation (Figure 1E, 1F, 2D, 3B, 3C, 4B, 4D, 4E, 5A, 5B, 5C, 5D, 6B, and Supplemental Figure S1E, S1F, S3B, S3C, S3F).

Error bars denote the standard deviation and ‘X’ or ‘o’ the mean (Figure 1F and 4C, and Supplemental Figure S1C).

### Modeling the relationship between power, elongation rate and size

We contextualized the observed relationships between elongation rates, spindle size, and breakage (Figure 4B, 5A, and 5B) in terms of a simple model of work energetics where Power = Work/Time. In this model, we assumed that the work required to bend and break the spindle is similar in the WT and the different mutant cases (except for *ase1Δ* cells), as they all produce a bent spindle of similar dimensions before breaking (Figure 5B). The variable times of elongation to reach the terminal size indicate that the power of the spindles in the mutant strains is proportional to the elongation rate. For instance, the 50% slower elongation rate in *klp9Δ* mutant suggests that it has 50% of the power of a WT spindle. Suppose we assumed that the power exerted by bent and straight spindles of a particular genotype are similar. In that case, this leads to a prediction that the ratio of elongation rates in bent versus straight spindles should be maintained even at different powers, consistent with our experimental measurements (red line, Figure 4C). These findings suggest that the consistent breakage point of the bent spindles is determined not by a timing mechanism but by the accumulation of mechanical strain in the bent spindle produced by spindle elongation.

### Microtubule number in the *S. pombe* spindle

Although the number and organization of MTs in different stages of fission yeast spindles have been estimated using the fluorescence intensity of tubulin-GFP (Loiodice et al., 2019) and cryo-EM (Ding et al., 1993; Ward et al., 2014). There is almost two-fold discrepancy in the literature on MT numbers in the fission yeast spindle. Some of this discrepancy arises from the fact that some of the data was collected from *cdc25* mutants, which exhibit larger spindles with increased numbers of MTs (Ding et al., 1993; Loiodice et al., 2019). The Ward data are based upon a wildtype haploid *S. pombe* strain but are only represented by two mid-sized spindles that appear to have fewer MTs than the average spindle, as determined by fluorescence intensity measurements in the same study. Especially as discrepancies in the absolute number of MTs and spindle size can make a very large difference in force estimates, additional data on the MT number at various stages of wildtype spindles are needed to arrive at a reliable measure of anaphase elongation forces.

### Source code

https://github.com/JoelLEMI/Analyses-of-bent-spindles-reveal-the-mechanics-of-anaphase-B-in-fission-yeast

## Acknowledgements

We thank present and past members of the Chang lab, Sophie Dumont, Pooja Suresh for valuable discussion. We thank Phong Tran for strains, and PomBase for valuable resources. This work was supported by grants to F.C. (NIH GM115185, NIH GM056836, NIH GM141796) and to F.C. and T.G.F. (NSF MCB 2213583)

## Abbreviations

CI: coefficient interval
DNA: deoxyribonucleic acid
EM: electron microscopy
ER: endoplasmic reticulum
GFP: green fluorescent protein
mCh: mCherry
MT: microtubule
mNG: mNeonGreen
NE: nuclear envelope
SPB: spindle pole body
WT: wild type

---

**Supplemental Table S1.**

| Number | Strain | Figure | Origin |
| --- | --- | --- | --- |
| FC2859 | *hta1-mCherry:natMX4 GFP-atb2:kanMX ade6- leu1-32 ura4-* | 1 and S1 | Lab collection |
| FC3241 | *mCherry-atb2:hphMX6 mis12-GFP:leu2+ leu1-32 ura4-D19 ade5-M21X* | 1, 2, 3, 4, 5, S1, S3 and S6 | Phong Tran |
| FC3374 | *mCherry-atb2:hphMX6 mis12-GFP:leu2+ ish1-GFP:kanMX* | 1 | This study |
| FC3246 | *klp9Δ:kanMX6 mCherry-atb2:hphMX6 mis12-GFP:leu2+ leu1-32 ura4-D19 ade5-M21X* | 2, 4 and 5 | Phong Tran |
| FC3247 | *h- cut7Δ:natMX6 pkl1Δ:ura4+ mCherry-atb2:HphMX6 mis12-GFP:leu2+ leu1-32 ura4-D19 ade5-M21X* | 2, 4 and 5 | Phong Tran |
| FC3248 | *h- klp2Δ:kanMX6 mCherry-atb2:HphMX6 mis12-GFP:leu2+ leu1-32 ura4-D19 ade5-M21X* | 2, 4 and 5 | Phong Tran |
| FC1984 | *ase1Δ:kanMX leu1-32::SV40-GFP-atb2[LEU1] leu1-32 ura4-D18 his7+* | 2, 4 and 5 | Lab collection |
| FC3245 | *h- klp9-GFP:kanMX mCherry-atb2:HphMX6 mis12-GFP:leu2+ leu1-32 ura4-D19 ade5-M21X* | 3 and S2 | Phong Tran |
| FC3375 | *ase1-mNeonGreen:KanMX mCherry-atb2:hphMX6* | 3, 6, 7, S3, S6 and S7 | Lab collection |

